# Photoperiod sensing of the circadian clock is controlled by *EARLY FLOWERING 3* and *GIGANTEA*

**DOI:** 10.1101/321794

**Authors:** Muhammad Usman Anwer, Amanda Davis, Seth Jon Davis, Marcel Quint

## Abstract

*ELF3* and *GI* are two important components of the Arabidopsis circadian clock. They are not only essential for the oscillator function but are also pivotal in mediating light inputs to the oscillator. Lack of either results in a defective oscillator causing severely compromised output pathways, such as photoperiodic flowering and hypocotyl elongation. Although single loss of function mutants of *ELF3* and *GI* have been well-studied, their genetic interaction remains unclear. We generated an *elf3 gi* double mutant to study their genetic relationship in clock-controlled growth and phase transition phenotypes. We found that *ELF3* and *GI* repress growth differentially during the night and the day, respectively. Circadian clock assays revealed that *ELF3* and *GI* are essential *Zeitnehmers* that enable the oscillator to synchronize the endogenous cellular mechanisms to external environmental signals. In their absence, the circadian oscillator fails to synchronize to the light-dark cycles even under diurnal conditions. Consequently, clock-mediated photoperiod-responsive growth and development is completely lost in plants lacking both genes, suggesting that ELF3 and GI together convey photoperiod sensing to the central oscillator. Since *ELF3* and *GI* are conserved across flowering plants and represent important breeding and domestication targets, our data highlight the possibility of developing photoperiod-insensitive crops by adjusting the allelic combination of these two key genes.

**One sentence summary:** ELF3 and GI are essential for circadian clock mediated photoperiod sensing.

## Introduction

Rotation of the earth around its axis results in rhythmic oscillations in light and temperature during a 24-hour day/night cycle. As a consequence of evolving under these predictable changes, organisms have developed internal timekeeping mechanisms known as the circadian clock that enable them to anticipate periodic changes in their surrounding environment (de Montaigu et al., 2010; Anwer and Davis, 2013). Circadian clocks consist of three pathways: inputs, core oscillators, and outputs. Input pathways deliver external cues (also known as *Zeitgeber*, German for time-givers), such as ambient light and temperature, to circadian oscillators. The timing information from the *Zeitgeber* is received by core-oscillator components known as *Zeitnehmer* (German for time-takers) that help to reset and synchronize the clock with the local environment (entrainment). Once entrained, the oscillators generate a ∼24h rhythmicity that can be sustained for long periods; even in the absence of environmental cues (i.e., free-running conditions, such as constant light and temperature conditions) (Inoue et al., 2017; Oakenfull and Davis, 2017). After synchronizing with the external environment, oscillators link to various processes to rhythmically regulate the levels of transcripts, proteins, and metabolites. This allows organisms to anticipate and adapt to the changing environment, such as seasonal changes in day length (photoperiod). The circadian clock thereby regulates various output pathways including photosynthesis, growth, disease resistance, starch metabolism, and flowering time (Andres and Coupland, 2012; Shin et al., 2013; Müller et al., 2014).

The central part of the clock, the oscillators, are composed of transcriptional-translational feedback loops (Nohales and Kay, 2016; Ronald and Davis, 2017). The *Arabidopsis thaliana* (Arabidopsis) oscillator consists of three such loops: a morning loop, an evening loop and a central oscillator. The central oscillator is comprised of two partially redundant myb-like transcription factors CIRCADIAN CLOCK ASSOCIATED 1 (CCA1) and LATE ELONGATED HYPOCOTYL (LHY), and a member of the PSEUDO-RESPONSE REGULATOR (PRR) family TIMING OF CAB EXPRESSION 1 (TOC1/PRR1). This is a dual negative feedback loop where respective morning and evening expression of *CCA1/LHY* and *TOC1* repress each other (Wang and Tobin, 1998; Alabadí et al., 2001; Huang et al., 2012). In the morning, the core-oscillator components CCA1/LHY repress *PRR7* and *PRR9,* which later repress *CCA1/LHY,* together constituting the morning loop (Zeilinger et al., 2006; Nakamichi et al., 2010; Adams et al., 2015; Kamioka et al., 2016). The evening expression of TOC1 represses *GIGENTEA (GI)*, which in turn activates *TOC1* and formulates the evening loop (Locke et al., 2006; Kim et al., 2007; Huang et al., 2012). Besides these three fundamental loops, a complex of three evening phased proteins (known as evening complex or EC), consisting of EARLY FLOWERING 4 (ELF4), ELF3 and LUX ARRYTHMO (LUX), has been identified as an essential part of the core oscillator (Nusinow et al., 2011; Herrero et al., 2012; Huang and Nusinow, 2016). The EC is connected to all three loops of the oscillator. By direct binding to their promoters, the EC represses the transcription of *PRR9* and GI (Helfer et al., 2011; Herrero et al., 2012; Mizuno et al., 2014; Ezer et al., 2017). A direct repression of *ELF3* by CCA1 connects the EC with the central oscillator (Lu et al., 2012; Kamioka et al., 2016).

ELF3 is one focus of this study and it encodes a multifunctional protein that regulates several physiological and developmental processes. Consistently, *elf3* null mutants display pleiotropic phenotypes such as long hypocotyl, accelerated flowering, elongated petioles, and arrhythmia under free-running conditions, suggesting that several important pathways are disrupted (Hicks et al., 2001; Kolmos et al., 2011; Herrero et al., 2012; Anwer et al., 2014; Box et al., 2014). In addition to its role as a member of the EC in the core oscillator, it functions as a *Zeitnehmer* in the light input pathway.

Therefore, plants lacking *ELF3* display severe light gating defects (McWatters et al., 2000). A physical interaction of ELF3 and PHYTOCHROME B (PhyB) establishes a direct link between the oscillator and photoreceptors (Liu et al., 2001; Kolmos et al., 2011). For the regulation of rhythmic growth, ELF3 mainly relies on the EC binding to the promoters of major growth regulators *PHYTOCHROME-INTERACTING FACTOR 4 (PIF4)* and *PIF5*, causing their transcriptional repression during the night (Nusinow et al., 2011; Raschke et al., 2015). However, ELF3 can also inhibit PIF4 by sequestering it from its targets (Nieto et al., 2014). Consistently, the lack of *PIF4/PIF5* repression in *elf3* mutants results in accelerated growth during the night (Nozue et al., 2007; Box et al., 2014). In addition to growth, *ELF3* controls flowering time by acting on the major floral activator FLOWERING LOCUS T (FT) via direct repression of *GI* (Reed et al 2000; Mizuno et al., 2014; Ezer et al., 2017). Interestingly, ELF3 repression of *FT* does not require *CONSTANS (CO)* (Kim et al., 2005). Further, under field conditions, ELF3 ensures a morning expression of *FT*, which is important for flowering under long days (Song et al 2018). Taken together, functional presence of *ELF3* is essential for both plant growth and development.

The second protein in the focus of this study is GI, a large, preferentially nuclear-localized protein with domains of unknown functions (Panigrahi and Mishra, 2015). The gene’s transcription is controlled by the circadian clock. Furthermore, it is post-transcriptionally regulated by light and dark (Fowler et al., 1999; David et al., 2006). GI regulates diverse developmental and physiological pathways. The role of GI in the control of photoperiodic flowering is well documented. Here, GI acts as a major activator of *FT* expression, either by directly binding to its promoter or by inducing the expression of *CO (Fornara et al., 2009; Sawa and Kay, 2011)*. Moreover, GI physically interacts with both red and blue light photoreceptors PhyB and ZEITLUPE (ZTL), respectively, indicating a functional role also in photomorphogenesis (Kim et al., 2007; Yeom et al., 2014). Consistently, *gi* mutants are defective in proper light responses and display elongated hypocotyls under both red and blue lights (Huq et al., 2000; Martin-Tryon et al., 2007). Although the underlying molecular mechanism of hypocotyl growth regulation is not fully understood, it relies at least partially on PIF4, since the growth promoting effect of *gi* mutations was fully masked by the absence of *PIF4 (de Montaigu et al., 2014; Fornara et al., 2015)*. The EC subunit *ELF4* is epistatic to *GI* in regulating hypocotyl length, suggesting that the *GI* effect on *PIF4* is EC dependent (Kim et al., 2012). However, *ELF4* masking of *GI* is specific to growth regulation because in flowering time control the genetic hierarchy between these two is reversed. Here, *GI* is epistatic to *ELF4*. To make the interaction between these two players even more interesting, both are working additively or synergistically in the control of the circadian clock (Kim et al., 2012). GI plays a pivotal role in generating robust circadian rhythms under natural conditions in a way that daily rhythms of its expression respond to day length that depends on the latitude of origin of Arabidopsis accessions (de Montaigu and Coupland, 2017).

Interestingly, GI co-localizes with the EC components ELF4, ELF3 and LUX in nuclear bodies (Yu et al., 2008; Herrero et al., 2012), where it physically interacts with ELF4 and ELF3 (Yu et al., 2008; Kim et al., 2013). ELF4 regulates GI subcellular localization and modulates its DNA binding ability by sequestering it from the nucleosome (Kim et al., 2013). Further, GI and ELF4 have differentially dominant influences on circadian physiological outputs at dusk and dawn, respectively (Kim et al., 2012). The functional importance of ELF3-GI interaction is unknown. However, it is reported that ELF3 regulates diurnal protein accumulation of GI by facilitating its degradation during darkness by a CONSTITUTIVE PHOTOMORPHOGENIC 1 (COP1) mediated proteasomal mechanism (Yu et al., 2008). Consistent with the finding that ELF3 binds to the *GI* promoter and represses its transcription (Mizuno et al., 2014), all components of the EC were found to bind the *GI* promoter in a CHIP-Seq experiment, demonstrating a direct relationship between *GI* and the EC (Ezer et al., 2017).

As mentioned above, the genetic hierarchy between *ELF4* and *GI* is relatively well understood (Kim et al., 2012). Based on the observations that mutations in EC components exhibit similar defects (Herrero et al., 2012), a conserved genetic relationship between GI and other EC components seems reasonable. On the other hand, the finding that ELF3 likely functions also independently of the EC (Nieto et al., 2014) opens the possibility for a different pattern of genetic interactions between *ELF3* and *GI*.

In this study, we provide genetic support for the biochemical evidence of an EC independent function of ELF3. We furthermore demonstrate that *ELF3* and *GI* are essential clock *Zeitnehmers* that are required to synchronize endogenous signals with the external environment. In their absence the circadian clock fails to properly respond to light signals, resulting in the breakdown of the photoperiod sensing mechanism. From an applied perspective, this interaction has the potential to generate photoperiod-independent crops, possibly allowing the cultivation of numerous day light sensitive species in currently non-permissive latitudes.

## Results

### *ELF3* and *GI* are essential for photoperiod responsive growth and development

*ELF3* and *GI* are two important factors involved in photoperiod responsive flowering (Andres and Coupland, 2012; Lu et al., 2012). A previous report has suggested that under long days (LD, 16h light/8 h dark) *GI* is epistatic to *ELF3* (Chou and Yang, 1999). *GI* is also epistatic to *ELF4*, another component of EC, further suggesting that flowering time control of the EC acts through *GI (Kim et al., 2012)*. However, it is unclear whether the suggested genetic hierarchy between *ELF3* and *GI* is universally applicable under a range of photoperiods. To investigate the environmental sensitivity of these genetic interactions in detail, we generated an *elf3-4 gi-158* double mutant (hereafter designated as *elf3 gi*) and measured flowering time in comparison to the corresponding single mutants *elf3-4* (hereafter designated as *elf3*) and *gi-158* (hereafter designated as *gi*), and the Ws-2 wild type (WT) under long day (LD, 16h light/8 h dark), short day (SD, 8/16), and neutral day (ND, 12/12) photoperiods. Consistent with reported phenotypes of *elf3* and *g*i null mutants (Zagotta et al., 1996; Fowler et al., 1999; Lu et al., 2012), *gi* and *elf3* single mutants flowered later and earlier, respectively, than WT under all photoperiods tested (Figure 1A). Furthermore, similarly to WT, both single mutant alleles flowered earlier in longer photoperiods than in shorter photoperiods, therefore displaying an intact response to the length of the light period. Interestingly, such a photoperiodic response was completely lost in the *elf3 gi* double mutant, where flowering time was unaffected by the photoperiod (Figure 1A). Moreover, while under LD and ND flowering time of *elf3 gi* was similar to *gi*, it was similar to *elf3* under SD (Figure 1A). Thus, unlike *ELF4*, where *GI* is epistatic under both LD and SD (Kim et al., 2012), no clear genetic hierarchy was observed between *ELF3* and *GI*, suggesting independent roles in flowering-time control.

**Figure 1.**
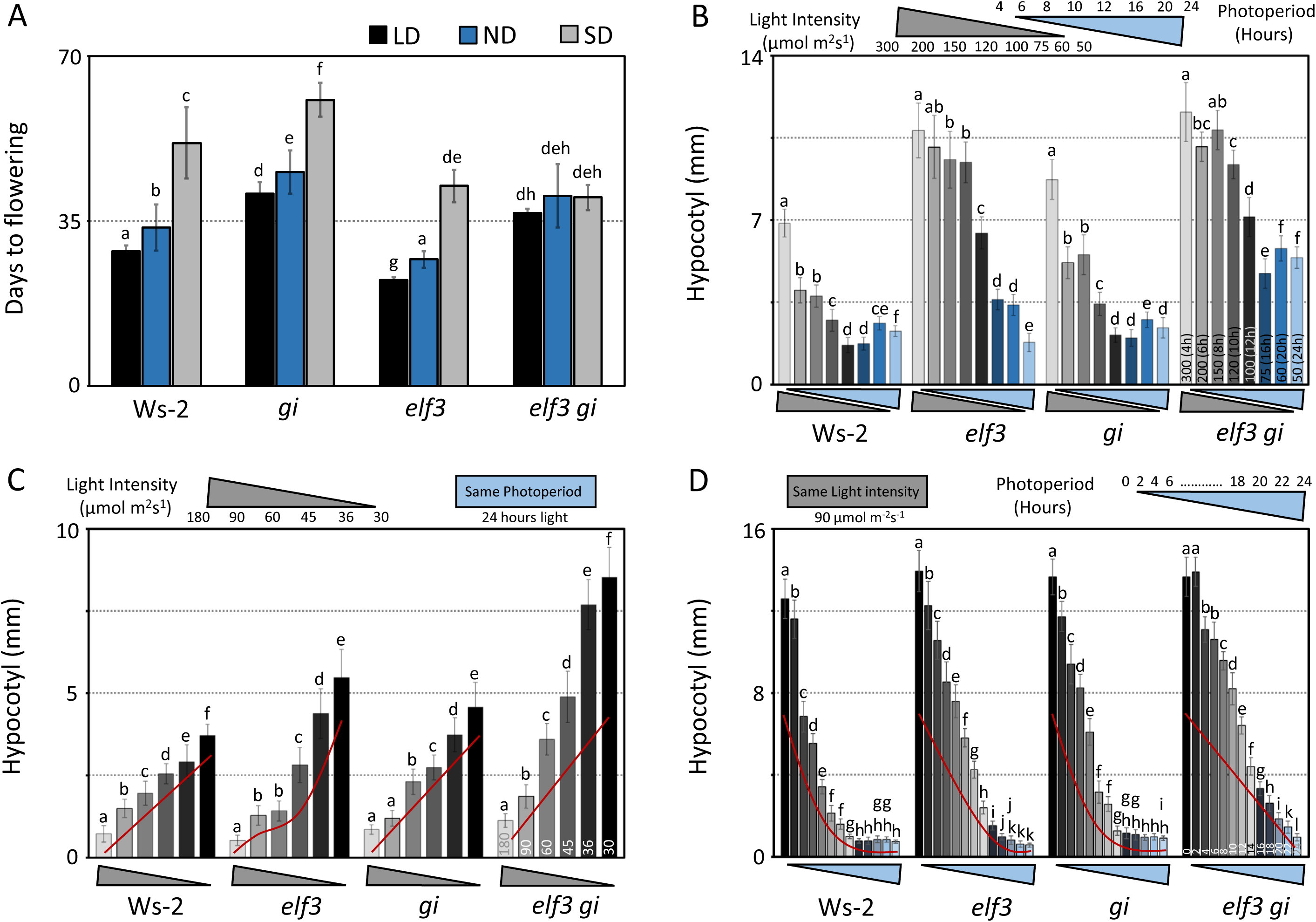
Photoperiod-responsive flowering and hypocotyl elongation require functional *ELF3* and *GI*. **(A)** Flowering time of Ws-2, *elf3*, *gi* and *elf3 gi* under LD (16h light: 8 h darkness), ND (12h light: 12h darkness), and SD (8h light: 16h darkness). Flowering time was counted as number of days to 1 cm bolt. Error bars represent standard deviation (StD); n≥24. Letters above the bars represent statistically significant differences calculated using one-way ANOVA (ANalysis Of VAriance) with *post-hoc* Tukey HSD (Honestly Significant Difference) Test, p<0.01. **(B)** Hypocotyl length of Ws-2, *elf3*, *gi* and *elf3 gi* under different photoperiods and light intensities. Length of the photoperiod and light intensity was set so that a total equal fluence (4320000 µmol) was achieved. Numbers at the bottom of the bars (*elf3 gi*) represent light intensities (µmol m^2^s^1^) and numbers within brackets represent photoperiods (hours). **(C)** Hypocotyl length of Ws-2, *elf3*, *gi* and *elf3 gi* under different light intensities. Numbers at the bottom of the bars (*elf3 gi*) represent light intensities (µmol m^2^s^1^). Red lines are trend lines used to better illustrate the data. **(B,C)** Hypocotyl lengths of Ws-2, *elf3*, *gi* and *elf3 gi* in darkness (DD) were: 18.88mm +− 1.46mm, 22.36mm +− 2.36mm, 19.40mm +− 1.90 and 21.29mm +− 2.34mm, respectively. **(D)** Hypocotyl length of Ws-2, *elf3*, *gi* and *elf3 gi* under different photoperiods (constant light: 90 µmol m^2^s^1^). Numbers at the bottom of the bars (*elf3 gi*) represent the length of the light period. For instance ‘8’ = 8h light: 16h darkness, and ‘12’ = 12h light: 12h darkness. Trend lines were used to better illustrate the data. Light intensity, 90 µmol m^−2^s^−1^. **(B,C,D)** Error bars represent standard deviation. n≥16. Letters above the bars represent statistically significant differences as described in (A). Seedling were grown for seven days under respective photoperiods and/or light intensities at constant 20°C before taking images.

Since transition from the vegetative to the reproductive phase is only one of several developmental processes influenced by the photoperiod, we next sought to determine whether *elf3 gi* is also insensitive to photoperiod during the early growth phase. A classic phenotypic output for vegetative growth is elongation of the juvenile stem (hypocotyl), which is determined by the length of the light period and the light intensity, so that an increase in photoperiod length or light intensity results in a decrease in hypocotyl growth. To examine if the hypocotyl elongation response to photoperiod and light intensity is intact in *elf3 gi*, we grew the *elf3 gi* double mutants along with WT and single mutants under varying photoperiods and light intensities, and then measured the length of the hypocotyl. In the first experiment, we varied both the light intensity and photoperiod in a way that under any given photoperiod-intensity setup, the plants received a total equal light fluence (4320000µmol) (Figure 1B). We found that in *g*i, despite longer hypocotyl under several photoperiod-intensity combinations, the overall pattern of the hypocotyl response to varying photoperiod-intensity was similar to WT (Figure 1B). Furthermore, the response pattern of *elf3* and *elf3 gi* to varying photoperiod-intensity were different than both the WT and *gi*. Specifically, as the photoperiod increased (light intensity decreased) we noted a much larger decrease in the hypocotyl lengths in the WT compared to *elf3* and *elf3 gi* (Figure 1B, 300 (4h) to 120 (10h)). These data suggested that photoperiod and/or light sensing in *elf3* and *elf3 gi* is compromised. To further investigate whether the observed growth anomalies were caused by a combination of defects in both light and photoperiod sensing pathways or a single pathway is a major contributor, we performed experiments where either the light intensity or photoperiod was changed while the other parameter was fixed. In such experiments we first measured hypocotyl length of the WT, *elf3*, *gi* and *elf3 gi* under different light intensities (constant light, 24 hrs photoperiod). We found that despite differences in the hypocotyl lengths of the *elf3*, *gi* and *elf3 gi* mutants under various light intensities, these mutants were fully capable of distinguishing differences in light intensities. This conclusion is based on the observation that, just like WT, an increase in the light intensity resulted in shortening of the hypocotyl length (Figure 1C), suggesting that the light signaling pathway is not disturbed in any of these mutants.

Next, we sought to determine the functional capacity of the photoperiod sensing pathway. In WT, the length of the photoperiod is inversely proportional to the length of the hypocotyl. However, this relationship is not linear. Until a critical photoperiod (14-16 h light) is reached, the growth inhibitory effect of the increased photoperiod remains intact. After this time point, a further increase in the photoperiod has almost no effect on growth (Niwa et al., 2009). To investigate the role of *ELF3* and *GI* in photoperiod growth control, we measured hypocotyl length of WT, *elf3*, *gi,* and *elf3 gi* seedlings grown under a range of photoperiods, from constant darkness (DD), with a gradual increase of 2 hours light periods, to 24 hours light (LL) (Figure 1D, Tables S1-S2, for better resolution see also Figure S1A-C). In confirmation of Niwa et al. (2009), an intact response to photoperiod was observed in WT with plants responding to an increase in day length with a decrease in hypocotyl length until the 16h photoperiod. After 16h, no significant decrease in hypocotyl length was observed. Albeit with an overall longer hypocotyl, WT-like response to the changing photoperiod was also observed in *gi* (Figure 1D, S1A-C, Tables S1-S2). Interestingly, both *elf3* and *elf3 gi* did not display an intact photoperiod response of growth inhibition. Unlike WT, the repressive action of longer photoperiods continued even after 16h. Notably, the effect of light repression was discontinued after 20h photoperiod in *elf3*, whereas, in *elf3 gi* it continued until LL (Figure 1D, S1A-C, Tables S1-S2). Thus, our data indicate a previously not recognized additive function of *ELF3* and *GI* in photoperiod sensing, which becomes obvious only in the absence of both genes.

The EC controls hypocotyl elongation by regulating the expression of *PIF4* (Nusinow et al., 2011). Under LD and SD, the length of the *elf4 gi* double mutant is similar to *elf4*, indicating that *ELF4* is epistatic to *GI (Kim et al., 2012)*. Since ELF3, like ELF4, is also a component of the EC, a similar genetic hierarchy could also be expected between *ELF3* and *GI*. If so, hypocotyl length of *elf3 gi* and *elf3* should be similar. However, we found that under both LD and SD *elf3 gi* was significantly longer than *elf3* (Figure S1A, Light periods 8 and 16), suggesting an additive function of *ELF3* and *GI*. Together, these data demonstrate that both *ELF3* and *GI* are essential for photoperiod sensing at both juvenile and adult stages of plant development. In addition, they provide genetic evidence for an EC-independent function of ELF3.

### *ELF3* and *GI* repress growth during night and day, respectively

Under diurnal conditions, the elongation of hypocotyl is gated by the circadian clock, allowing maximum growth to occur at dawn or early morning under SD and LD, respectively (Nozue et al., 2007). By repressing growth during the night, *ELF3* functions as an important factor in clock gating. Consistently, *elf3* mutants have been reported to lose the normal gating response, resulting in maximum growth during the night (Nozue et al., 2007; Box et al., 2014). The role of *GI* in clock-controlled growth, however, remains largely unknown. The additive growth phenotype of *elf3 gi* (Figure 1C-D, S1A), reveals two possibilities: first, both *ELF3* and *GI* work cooperatively at a similar time of day. If so, the loss of both in the *elf3 gi* double mutant results in an increased growth at that particular time. Alternatively, both repress growth at a different time of the day-night cycle, resulting in an enhanced growth in *elf3 g*i at separate times. To dissect these possibilities, we measured growth rates of WT, *elf3*, *gi* and *elf3 gi* every hour for two days under SD and LD using infrared imaging, which allowed growth monitoring also in darkness (Figure 2A-H). As reported previously (Nozue et al., 2007), maximum growth in WT was observed during the early morning (SD at ZT0 and LD at around ZT4 (Figure 2A,E). In *elf3*, growth rates were increased with maximum elongation detected during the night under both SD and LD (Figure 2B,F, and Table S3), confirming the night-specific repressive function of ELF3 in elongation growth. The *gi* mutant displayed a broader growth peak during early morning under SD, and afternoon under LD (Figure 2C,G, and Table S3). In *elf3 gi*, growth was pronounced during the night. However, in contrast to WT and both single mutants, growth rates did not peak at a specific time of day, but instead remained on a rather constant level. Compared to WT and the single mutants, the rate of elongation growth was increased during both day and night (Figure 2D,H, and Table S3). Taken together, while we can confirm the previously described growth-repressive role of ELF3 during the night, our results reveal an unknown role of GI in repressing growth specifically during day times. For effective gating of clock-controlled growth, both ELF3 and GI are essential.

**Figure 2.**
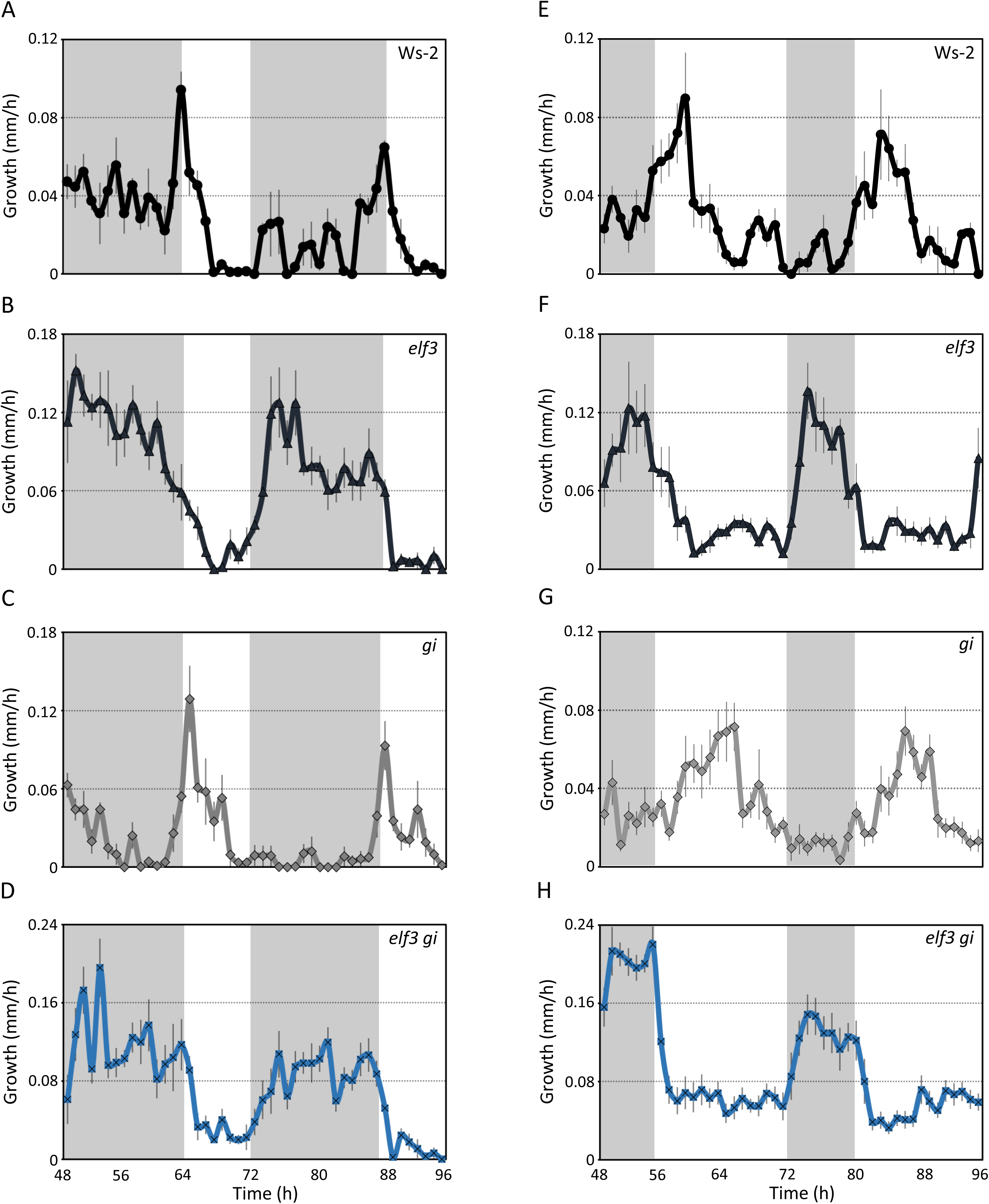
*ELF3* and *GI* repress growth during night and day, respectively. **(A-H)** Growth rate of Ws-2, *elf3*, *gi* and *elf3 gi* under SD (A-D) and LD (E-H). Starting from the third day, photographs were taken every hour using a modified Infra-red camera. Non-shaded areas in the graph represent daytime, shaded areas represent nighttime. Error bars represent standard error of the mean (S.E.M.); n≥8. The experiment was repeated three times with similar results. Light intensities were: SD 50 µmol m^2^s^1^ and LD 30 µmol m^2^s^1^.

### The circadian oscillator is dysfunctional in *elf3 gi* double mutants

Since *ELF3* and *GI* are important components of the circadian clock (Mizoguchi et al., 2005; Anwer et al., 2014), we asked whether the photoperiod insensitivity of *elf3 gi*, as revealed by growth and flowering behavior shown above, could be attributed to a malfunctional oscillator. To investigate the interactive role of *ELF3* and *GI* in the clock, we monitored the expression of the *CCR2:LUC* reporter under constant light (LL) in WT, *elf3*, *gi* and *elf3 gi* plants that were previously entrained under LD, ND or SD (Figure S2 A-C). As expected for a functional oscillator, WT displayed a robust rhythm. In contrast, no rhythmic expression of the reporter was detected in *elf3* and *elf3 gi*. The *gi* mutant was also rhythmic albeit with lower amplitude (Figure S2A-C). Moreover, the levels of *CCR2:LUC* in *elf3 gi* were higher than the WT, and the single mutants *elf3 and gi* (Figure S2-C), possibly hinting to an additive function of *ELF3* and *GI* in the clock.

Using the same data, we next calculated the free-running period of the aforementioned lines. Irrespective of the photoperiod provided for entrainment, we found that the WT displayed a similar free-running period (Figure S2D). Compared to WT, an acceleration in clock speed was observed in *gi* (Figure S2D). Like WT, the photoperiod used during entrainment had no effect on *gi* periodicity (Figure S2D). Consistent with their arrhythmic phenotypes, no regular pattern of periodicity response to photoperiod was detected in *elf3* and *elf3 gi*, which is why these lines had to be excluded from this particular analysis.

Next we assessed the precision of the oscillator by calculating the relative amplitude error (RAE). An RAE value of “0” represents a perfect rhythm, whereas an RAE of “1” typifies no rhythm (Anwer et al., 2014). A general cutoff value of 0.5 is normally used to distinguish between a robust and a dysfunctional oscillator. As expected for a fully functional clock, the WT displayed a very low RAE after all entrainments (Figure S2E). The RAE measured for *gi* was significantly higher than the WT but lower than 0.5, suggesting a compromised but functional clock (Figure S2E). Consistent with their arrhythmic phenotype, the RAEs of *elf3* and *elf3 gi* were extremely high (RAE>0.6), indicating a dysfunctional oscillator. Collectively, these data suggest that, irrespective of the functional ability of the oscillator, the photoperiod during preceding entrainment does not significantly affect important clock parameters such as period and precision.

### Clock entrainment to light signals requires both a functional *ELF3* and *GI*

Several clock mutants that are arrhythmic under free-running conditions, display robust oscillations under diurnal conditions, suggesting that the oscillator is still capable of reacting to persistent environmental changes (Yamashino et al., 2008). The complete lack of response of the *elf3 gi* double mutant to photoperiod (Figure 1A-D and S1A-C), however, prompted us to think otherwise. Specifically, we hypothesized that the oscillator in *elf3 gi* might not be responsive to light signals even under diurnal conditions. To test this hypothesis, we monitored the expression of major central-oscillator genes *CCA1*, *TOC1*, *PRR9*, and *GI* under diurnal conditions with SD, ND and LD light cycles (Figure 3A-H, Figure S3A-D). In WT, the expression profiles of all these genes were consistent with previous data (Kolmos et al., 2011; Anwer et al., 2014), with *CCA1* and *PRR9* peaking in the morning, and *TOC1* and *GI* peaking in the evening (Figure 3A-H). In *gi*, the expression of *TOC1* was lower than the WT under SD and LD, and was higher under ND. Consistent with previous reports for *gi* null mutants (Fowler et al., 1999; Kim et al., 2012) the levels of *PRR9* and *CCA1* were lower than WT under all light cycles except ND where *gi* and WT displayed similar *CCA1* expression (Figure 3A-H, S3A-C). In agreement with published data, expression of *TOC1*, *PRR9* and *GI* in *elf3* was higher than in WT, while *CCA1* expression was lower (Hall et al., 2003; Kolmos et al., 2011; Anwer et al., 2014) (Figure 3A-H, S3A-C). Importantly, in both *elf3* and *gi*, albeit differences in expression levels, the overall shape of the expression patterns of all genes tested was similar to WT (Figure 3A-H, S3A-D). These data thus indicate an aberrant but functional oscillator in *elf3* and *gi* single mutants, which is capable of responding to environmental signals and generating robust rhythms under diurnal conditions. In *elf3 gi* double mutants, however, besides *PRR9* under LD, no detectable response to diurnal light signals were observed (Figure 3A-H, S3A-D). The expression profile of all clock genes tested were different from both single mutants and WT. Most importantly, the characteristic peaks of expression of these genes, clearly detectable in WT, *elf3* and *gi,* were absent in *elf3 gi*. Most of the genes displayed a constant higher or lower expression, which was irresponsive to changes in the light during a diurnal cycle (Figure 3A-H, S3A-D). These data demonstrate that only in the absence of both *ELF3* and *GI,* the circadian oscillator is insensitive to persistent light-input cues. Thus, *ELF3* and *GI* are essential *Zeitnehmers* that are required for clock entrainment to external light cycles.

**Figure 3.**
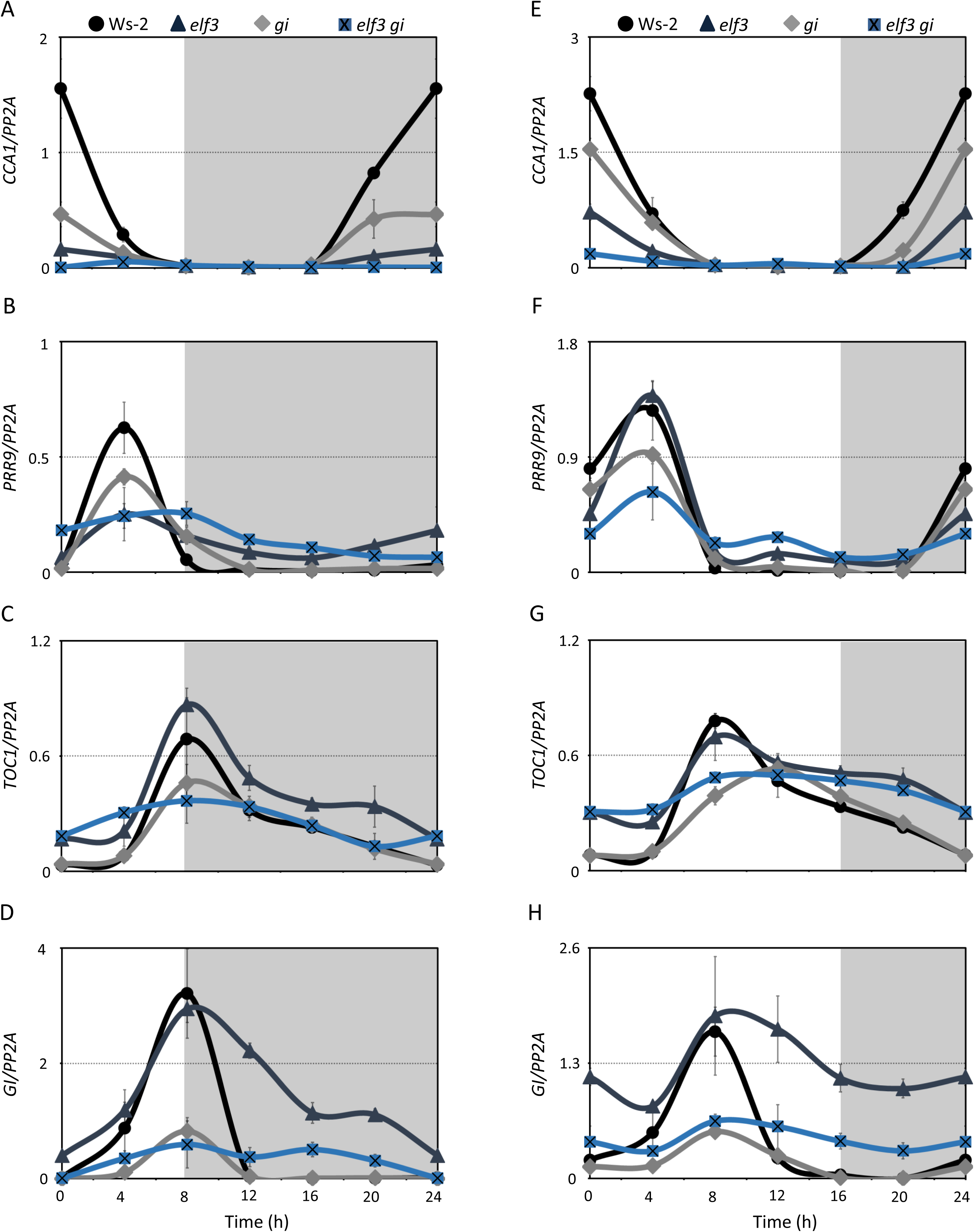
*ELF3* and *GI* are required for clock entrainment. Transcript accumulation of different clock oscillator genes *CCA1* **(A,E)**, *TOC1* **(B,F),** *PRR9* **(C,G)**, and *GI* **(D,H)** in Ws-2, *elf3*, *gi* and *elf3 gi* under SD (left panel (A-D)) and LD (right panel (E-H)). Error bars represent the standard deviation of three biological replicates. Expression levels were normalized to *PROTEIN 19 PHOSPHATASE 2a subunit A3* (*PP2A*). Non-shaded areas in the graph represent time in LL, shaded areas represent time in DD.

### *ELF3* and *GI* are essential to establish endogenous and light signaling links

Once entrained, the circadian clock regulates several key endogenous processes such as gene expression and ensures their precise synchronization with the external environment. This internal-external signaling synchronization is vital for several clock-controlled pathways such as flowering time and hypocotyl elongation. Since the oscillator in *elf3 gi* failed to establish a link with the external light signals, such a synchronization could potentially be lost in *elf3 gi*, explaining its photoperiod-insensitive flowering and growth. To test this hypothesis, we monitored diurnal expression behaviour of key clock-regulated genes involved in photoperiod-responsive flowering and hypocotyl elongation as a proxy.

To investigate the functional ability of the *elf3 gi* oscillator to regulate its target genes, we first monitored the expression of the key flowering-time genes *GI*, *CO* and *FT* under SD, LD and ND (Figure 4, S4 A-C). Consistent with previous reports, we detected a rhythmic expression of *GI*, *CO* and *FT* in WT (Fowler et al., 1999; Sawa and Kay, 2011), with *GI* expressed during the day with peak levels at ZT8, *CO* showing dual peaks, a smaller one at ZT8 and another one at ZT16-20. The maximum levels of *FT* were detected at dusk at ZT 8, ZT12 and ZT 16 for SD, ND and LD, respectively. (Figure 4, S4A-C). Consistent with the late flowering phenotype of the *gi* null mutant, the expression of *CO* and *FT* was barely detectable in *gi (*Figure 4B,C,F,G and S4B,C). In *elf3,* the expression of *GI* was higher at almost all time points (Figure 4A,E and S4A), consistent with the direct repression of *GI* by ELF3 (Mizuno et al., 2014; Ezer et al., 2017) as well as the early flowering phenotype of *elf3-4*. The expression of *CO* and *FT* was also much higher at several timepoints during the day and night (Figure 4B,C,F,G and S4B,C). The expression pattern of *CO* and *FT* in *elf3 gi* was similar to *gi*. Notably, no diurnal peak of expression was observed in the *elf3 gi* double mutant for any of the genes tested, with the overall expression hardly fluctuating over the entire diurnal cycle (Figure 4, S4A-C), reaffirming a dysfunctional oscillator.

**Figure 4.**
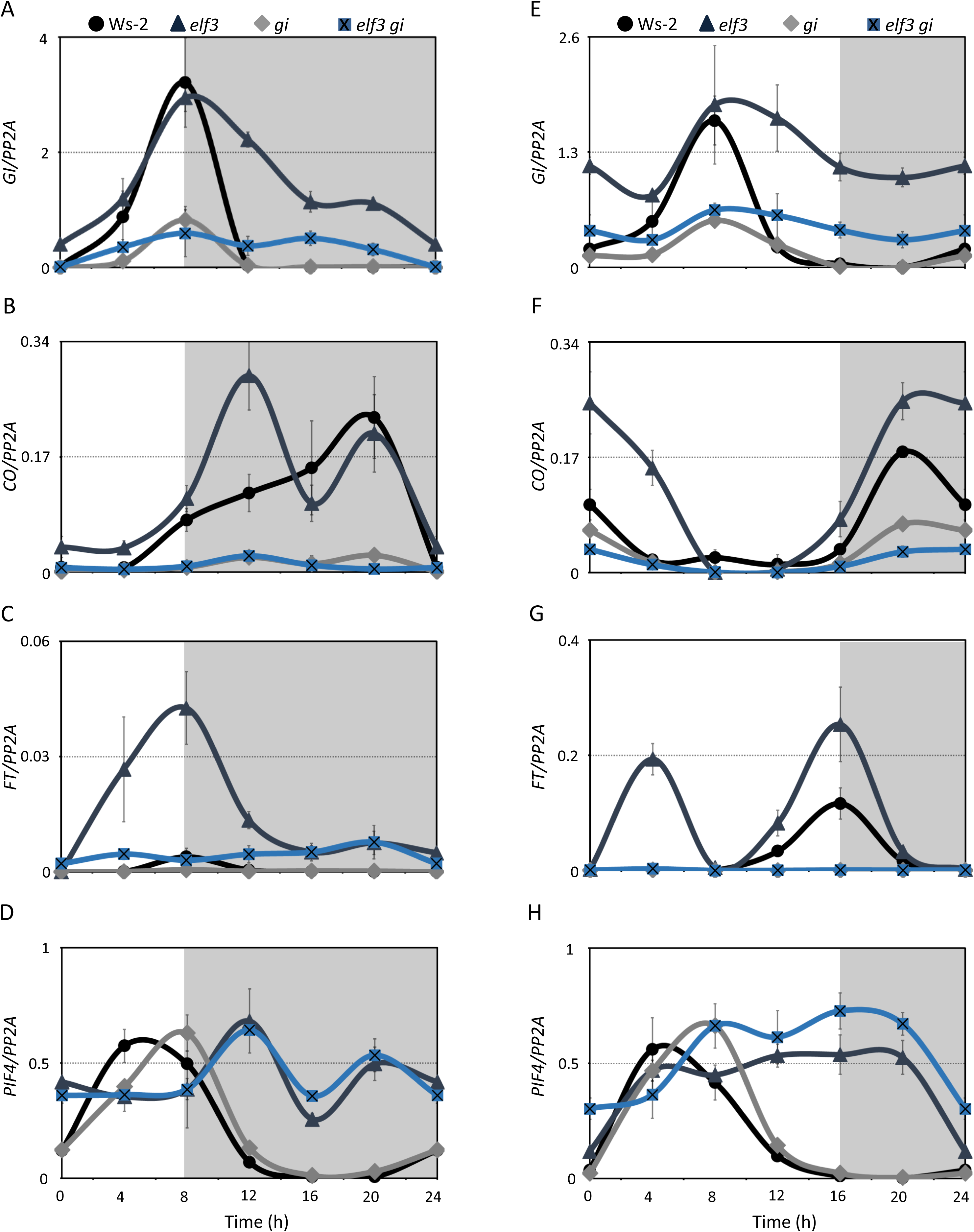
Endogenous and environmental signal synchronization requires functional *ELF3* and *GI*. Transcript accumulation of different flowering time genes *GI* **(A,E)**, *CO* **(B,F)**, *FT* **(C,G)**, and major growth promoter *PIF4* **(D,H)** in Ws-2, *elf3*, *gi* and *elf3 gi* under SD (left panel (A-D)) and LD (right panel (E-H)). Error bars represent the standard deviation of three biological replicates. Expression levels were normalized to *PROTEIN 19 PHOSPHATASE 2a subunit A3* (*PP2A*). Non-shaded areas in the graph represent time in LL, shaded areas represent time in DD.

We further validated these results by monitoring the expression of the major growth promoter *PIF4* under same conditions as mentioned above (Figure 4D,H and S4D). Consistent with their long hypocotyls, an overall higher expression of *PIF4* was observed in *elf3* and elf 3 *gi* under all conditions tested. Furthermore, in *gi, PIF4* followed a similar clock-regulated diurnal pattern as that of WT (Figure 4D,H and S4D). Under ND, *PIF4* expression in *elf3* also followed a diurnal pattern. However, it showed a characteristic light regulated profile with a gradual decrease in expression during the light period and a gradual increase during the dark period (Figure S4D). Interestingly, under ND, the *PIF4* expression in the *elf3 gi* double mutant was completely different from the diurnal patterns in WT and single mutants. Compared to WT, the level of *PIF4* was higher in *elf3 gi* at almost all time points, explaining for example its extreme growth phenotype shown in Figure 1. Further, *elf3 gi* displayed neither the clock regulated *PIF4* profile as observed for *gi*, nor the light regulated expression as observed in *elf3* (Figure S4D). Under SD and LD, albeit marginal but consistent higher levels of *PIF4* in *elf3 gi* under LD, overall similar expression patterns were observed in *elf3* and *elf3 gi* (Figure 4D,H). A closer examination revealed that with the exception of few points (LD: ZT0, ZT4 and ND: ZT16) the *PIF4* levels *in elf3 gi* remained almost similar throughout the diurnal cycles (Figure 4D,H and S4D). Collectively, these data demonstrate that both *ELF3* and *GI* are required for clock entrainment and thereby for the generation of rhythmic endogenous processes synchronized with the external signals.

## Discussion

The circadian clock is an important time keeping mechanism that synchronizes internal cellular mechanisms to the external environment. Light is the primary cue that provides timing information to the clock (Inoue et al., 2017; Oakenfull and Davis, 2017). While light sensing by the photoreceptors is well understood, it remains unclear how this information is perceived by the central oscillator. Here, we show that clock components *ELF3* and *GI* are essential to perceive light input into the clock and thereby for the measurement of the photoperiod. Absence of these components results in a dysfunctional oscillator, even under diurnal conditions, failing to regulate photoperiod-responsive growth and development.

Single loss of function mutants of individual EC components exhibit similar clock, hypocotyl and flowering time phenotypes, indicating that they work cooperatively (Nusinow et al., 2011; Herrero et al., 2012). Recent biochemical data has suggested that ELF3 can also function independently of the EC (Nieto et al., 2014). However, conclusive genetic evidence supporting the biochemical data is lacking. A previous study reported a clear genetic hierarchy between *ELF4* and *GI* with *ELF4* being epistatic to *GI* in control of hypocotyl elongation. *Vice versa*, *GI* is epistatic to *ELF4* in flowering time regulation (Kim et al., 2012). In our study, we did not observe such genetic relationships for *ELF3* and *GI*. Taking into account that ELF3 and ELF4 function together in the EC (Nusinow et al., 2011; Herrero et al., 2012; Quint et al 2016), this is somewhat surprising, supporting the proposed EC independent function for ELF3 (Nieto et al., 2014). The phenotypes we observed in single and double mutants for hypocotyl elongation suggest an additive function of *ELF3* and *GI* in controlling elongation growth (Figure 1C-D, S1A-C), whereas in flowering time regulation *ELF3* and *GI* were epistatic to each other under SD and LD, respectively (Figure 1A). The observation that the levels of key flowering time regulator *FT* are almost identical in both *gi* and *elf3 gi* under LD, but are slightly and consistently higher in *elf3 gi* under SD (Figure 4C,G) might explain the early flowering of the *elf3 gi* under SD (Figure 1A). However, whether these minute differences in *FT* expression are indeed sufficient for the early induction of flowering in *elf3 gi* under SD is questionable. In circadian clock control, *elf3 gi* displayed similar additive/synergistic phenotypes (Figure 3A-H) as reported for *ELF4* and *GI*. Collectively and in agreement with the biochemical data, our genetic analyses demonstrate that *ELF3* function is not solely dependent on the EC.

The molecular mechanism by which GI controls growth is not fully understood. An elongated hypocotyl of *gi* mutants under red and blue light suggested a repressive role in photoreceptor mediated growth inhibition (Huq et al., 2000; Martin-Tryon et al., 2007). Recent data demonstrated that GI requires PIF4 for growth regulation (de Montaigu et al., 2014; Fornara et al., 2015). Since the EC regulates *PIF4 (Nusinow et al., 2011)* and *ELF4* is epistatic to *GI (Kim et al., 2012)*, a role of *GI* upstream of the EC in growth regulation has been proposed (de Montaigu et al., 2014). However, our data, especially an additive hypocotyl phenotype and increased levels of *PIF4* in *elf3 gi* (Figure 1C-D, 4H, S1A-C, S4D), advocate an independent repressive action of GI on *PIF4*.

As growth and developmental phenotypes investigated in this study depend on the circadian clock, we asked whether ELF3’s and GI’s function in the clock might be able to explain the observed effects. By a “gating” mechanism the clock ensures that maximum growth happens at the correct time of day. In WT, under SD and LD growth rates peak in the early morning coinciding with the maximum expression of *PIF4 (Nozue et al., 2007)*. To coordinate this timing of growth rates, TOC1 and EC components including ELF3 repress growth during the late-evening and night, respectively (Nozue et al., 2007; Box et al., 2014; Zhu et al., 2016). In this study, we demonstrate that *GI* is also essential for clock mediated gating. GI represses growth during mid-day to late afternoon, thereby contributing to restricting growth peaks to the morning, resulting in normal rhythmic growth. Consistently, the loss of day and night time gating response in *elf3 gi* double mutants results in uncontrolled elongation growth (Figure 2D,H). Based on these observations we propose a model of rhythmic growth incorporating *ELF3* and *GI*. In this model, ELF3 and GI gate growth mainly by repressing *PIF4* during the night and late afternoon, respectively, allowing it to accumulate only during the early morning under LD. The morning accumulation of *PIF4* induces its downstream targets that consequently trigger cellular growth (Figure S4E).

The gating properties of the circadian clock are mainly dependent on its ability to synchronize internal cellular mechanisms with the external environment. Although after entrainment the clock maintains the same rhythm in the absence of the external input, in nature these free-running conditions almost never exist. Thus, proper clock responses to consistent external cues during a diurnal cycle are crucial for the synchronization of endogenous and environmental signals. Interestingly, arrhythmic clock genotypes, such as null mutants of the EC members *ELF3*, *ELF4* and *LUX*, as well as overexpressors of *CCA1* and *TOC1,* exhibit a non-functional oscillator under free-running conditions, but they are fully capable of generating robust rhythms under diurnal conditions (Fowler et al., 1999; Makino et al., 2002; Hall et al., 2003; Kolmos et al., 2011; Kim et al., 2012). Even higher order clock mutants including *cca1-1 lhy-11 toc1-2,* which lack the entire central oscillator, can generate rhythms under cycling conditions (Yamashino et al., 2008). The data presented in this study demonstrate that the absence of the two components *ELF3* and *GI* is sufficient to make the oscillator arrhythmic under both free-running and even under diurnal conditions (Figure 3A-H, S3A-D). We demonstrated that ELF3 and GI serve as important *Zeitnehmers* that are essential for clock entertainment. In their absence, the oscillator cannot effectively perceive external timing cues provided by cycling light conditions and, thus, fails to generate robust rhythmic oscillations of the downstream endogenous outputs. A closer look at the transcriptional profile of the major core-oscillator genes and the clock-regulated output genes under diurnal conditions in *elf3 gi* suggests that the entire clock-regulated transcriptome seems arrested (Figure 3A-H, S3A-D). As such, even changes in the environmental conditions during a diurnal cycle had almost no effect on the oscillator and were unable to fully release the clock-regulated transcriptome from its arrested state (Figure 4A-H, S4A-D). This should rationally lead to a breakdown of any clock-control output pathway. Consistently, photoperiod-responsive flowering and growth was disrupted in *elf3 gi* (Figure 1A-d, S1A-C). Notably, light regulated processes that are independent of the circadian clock seem to be intact in *elf3 gi*. A continuous inhibition of hypocotyl length under increasing photoperiod and light intensities (Figure 1C-D, Tables S1-S2) along with marked differences in growth rate during the light and dark phase in *elf3 gi* support this notion (Figure 2D,H and Table S3). Taken together, our data demonstrate that *ELF3* and *GI* control the circadian clock Zeitgeber-Zeitnehmer interface, enabling the oscillator to synchronize internal cellular mechanisms to the external environment.

Orthologues of *ELF3* and *GI* have been identified in several higher plants. Both genes have been prime breeding targets in crops for flowering time (Faure et al., 2012; Bendix et al., 2015; Panigrahi and Mishra, 2015; Huang and Nusinow, 2016). The *elf3 gi* double mutants develop rather normally and flower at the same time irrespective of the photoperiod. If similar genetic and functional relationships between *ELF3* and *GI* exist in economically important crops as reported here for Arabidopsis, breeders could develop photoperiod-insensitive varieties lacking *ELF3* and *GI* that would be independent of latitudinal photoperiodicity (Soyk et al., 2016).

## Materials and methods

### Plant material

All genotypes used were in the Ws-2 genetic background. The *elf3-4* null mutant (Liu et al., 2001) was previously described in (Zagotta et al., 1996; Hicks et al., 2001). The *gi-158* mutant was obtained by us in an ENU (*N*-ethyl-*N*-nitrosourea) genetic screen and will be explained in detail elsewhere. The *gi-158* is likely a null mutant that contains a premature stop codon resulting in a truncated protein of 146/1173 amino acids (Figure S5A-B). The flowering time and hypocotyl phenotypes of *gi-158* were very similar to the *gi-11* null mutant (Fowler et al., 1999) (Figure S5A-B). The double mutant *elf3-4 gi-158* was generated by crossing *elf3-4* and *gi-158,* and was confirmed by genotyping (Figure S5C). Marker used for genotyping were: *gi-158*, (forward ACTCATTACAACCGTCCCATCTA, reverse, GCGCATGAACACATAGAAGC (XbaI) *elf3-4* (forward TGCAGATAAAGGAGGGCCTA, reverse, ATGGTCCAGGATGAACCAAA. The *CCR2:LUC* line was used as previously described (Doyle et al 2002).

### Growth conditions

For luciferase assays, seeds were surface-sterilized and plated on MS medium containing 3% sucrose. Following ∼3 days stratification at 4°C, seedlings were entrained for 7 days, either under LD, ND, or SD cycles (∼100 µmol m-^2^s^−1^) with constant temperature of 20°C. The bioluminescence measurement and data analysis was performed as described (Hanano et al., 2008). For hypocotyl assays, seedlings were grown on *Arabidopsis thaliana* solution medium, as described previously (Lincoln et al., 1990). Hypocotyl length was determined for seedlings grown under varying photoperiod or light intensities for 7 days. White fluorescent lamps (T5, 4000 K) were used as light source. Seedlings were imaged, and hypocotyl elongation was measured using the *Rootdetection* program (http://www.labutils.de/rd.html). For flowering time measurement, plants were grown on soil containing a 3:1 mixture of substrate and vermiculite in phytochambers with LD, ND, SD cycle (white fluorescent light: 90 µmol m^−2^s^−1^, constant 22°C). Flowering time was scored at the time of bolting (1 cm above rosette leaves) as the total number of days to bolt. For all experiments, data loggers were used to monitor growth conditions.

### Infra-red photography for growth rate measurement

Seedlings were grown as described above with the following exception: to facilitate imaging unobstructed in air, seedlings were grown vertically on an agar ledge formed by removing part of agar in a square petri plate. Seeds were placed in small ridges on top of the agar. Imaging was started as soon as the cotyledons emerged from the seed coat. Photographs were taken every 60 minutes for 48 hours in LD cycles (white fluorescent light: 30 µmol m^−2^s^−1^, constant 20°C) or SD cycles (white fluorescent light: 50 µmol m^−2^s^−1^, constant 20°C). To image growth in day-night cycles we built an infrared imaging platform consisting of a modified camera with IR long pass 830 nm cut filters (Panasonic G5). Illumination was achieved using 880 nm IR backlights (Kingbright BL0106-15-29). Image stacks were analyzed using ImageJ (http://rsb.info.nih.gov/ij). Data loggers were used to monitor the growth conditions.

### Expression Analysis

Total RNA was isolated with the NucleoSpin® RNA Plant kit (Macherey-Nagel) following the manufacturer’s protocol from 1-week-old seedlings entrained under SD, ND or LD (90 µmol m^−2^s^−1^, constant 20°C). Quantitative RT-PCR and primer sequences were previously described (Kolmos et al., 2009) with following modifications: ABsolute Blue qPCR SYBR Green (ThermoFisher®) was used instead of iQ SYBR Green (Biorad). Agilent Mx3005P or AriaMx realtime system (Agilent®) were used instead of BioRad. Data loggers were used to monitor growth conditions.

## Supplementary Figures

**Figure S1.**
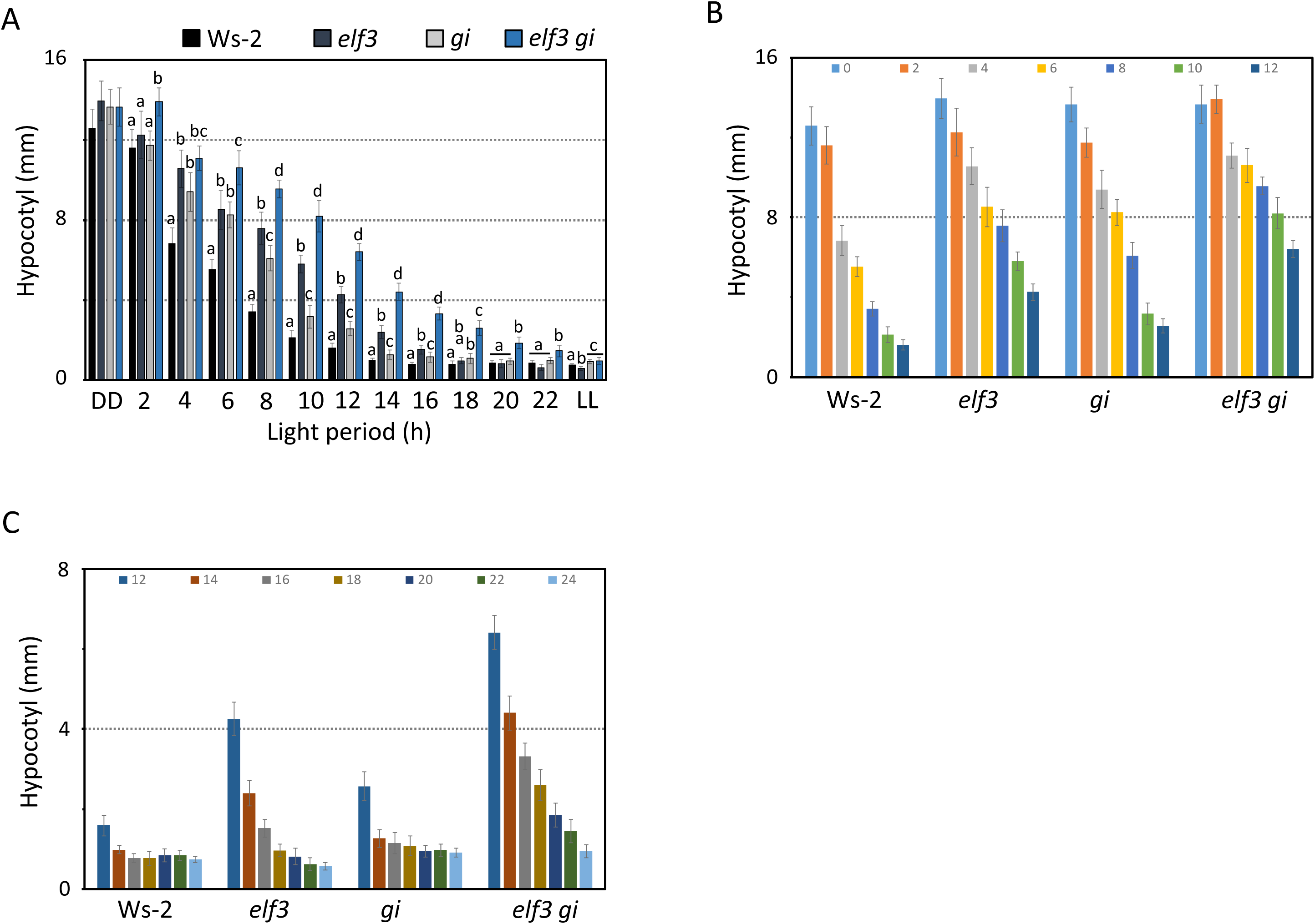
Effect of photoperiod on hypocotyl elongation. Related to Figure 1. **(A-C)** Hypocotyl length of Ws-2, *elf3*, *gi* and *elf3 gi* under different photoperiods as shown in Figure 1D. **(B-C)** For clarification, data shown in Figure 1D is split into two photoperiod ranges 0-12 **(B)** and 12-24 **(C)**. Growth condition, error bars and statistical analysis as described in Figure 1B, except the letters above bars represent differences among genotypes.

**Figure S2.**
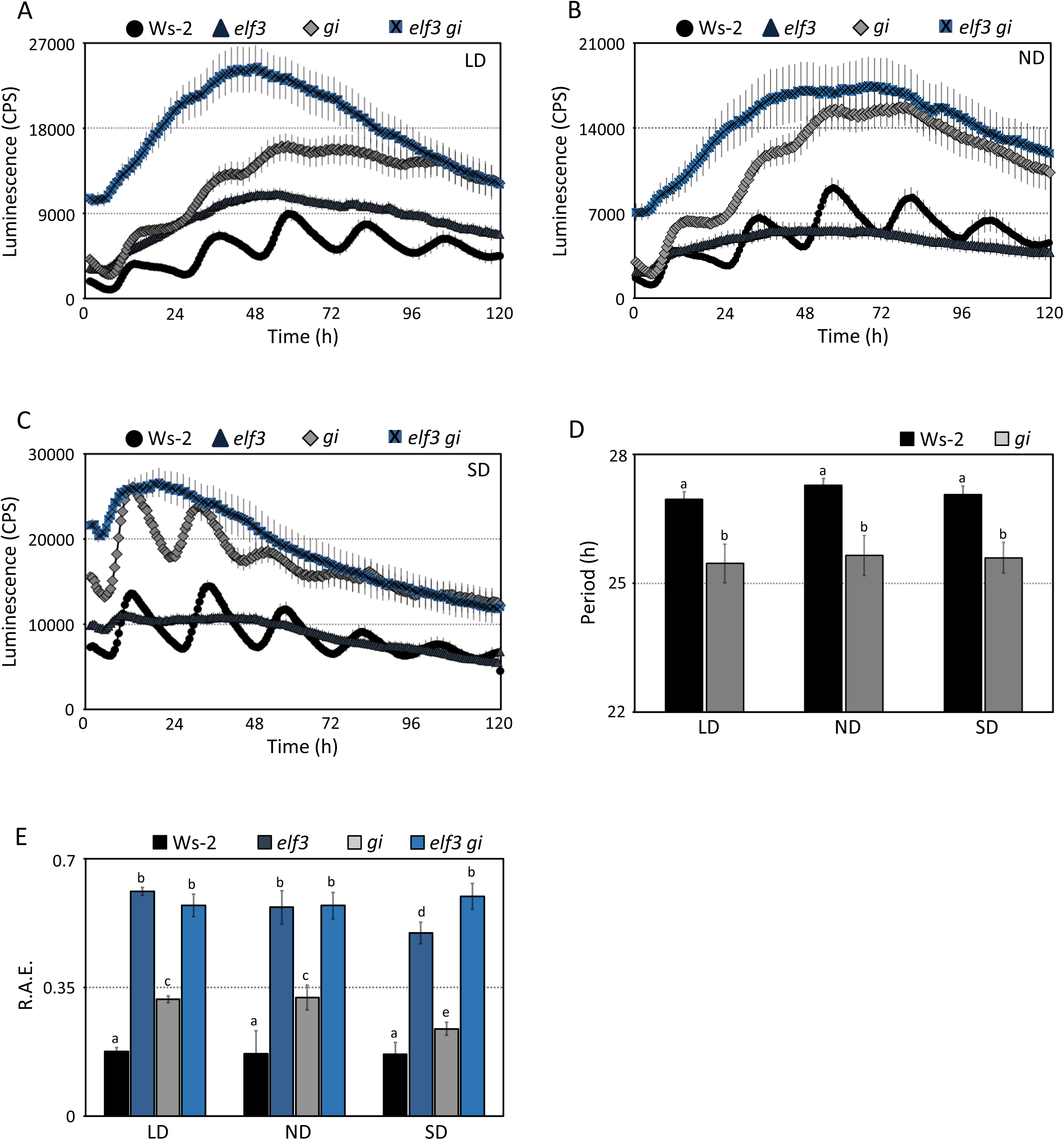
Oscillator functionality is independent of the preceding entrainment. **(A-C)** Free-running profile of *CCR2::LUC* expression in Ws-2, *elf3*, *gi* and *elf3 gi* under LD (A), ND (B) and SD (C). The plants were entrained for 7 days under respective light dark cycles, followed by transfer to LL and measurement of *CCR2::LUC* expression for 5 days. Error bars represent SEM and are shown on every third reading. **(D)** Period and **(E)** Relative amplitude error (R.A.E.) estimates after entrainment under different photoperiods as shown in (A-C). Error bars are SEM; n=48. Significance as described in Figure 1. Because of the arrhythmic nature of the *elf3* and *elf3 gi*, these lines were excluded from period analysis in **(D)**.

**Figure S3.**
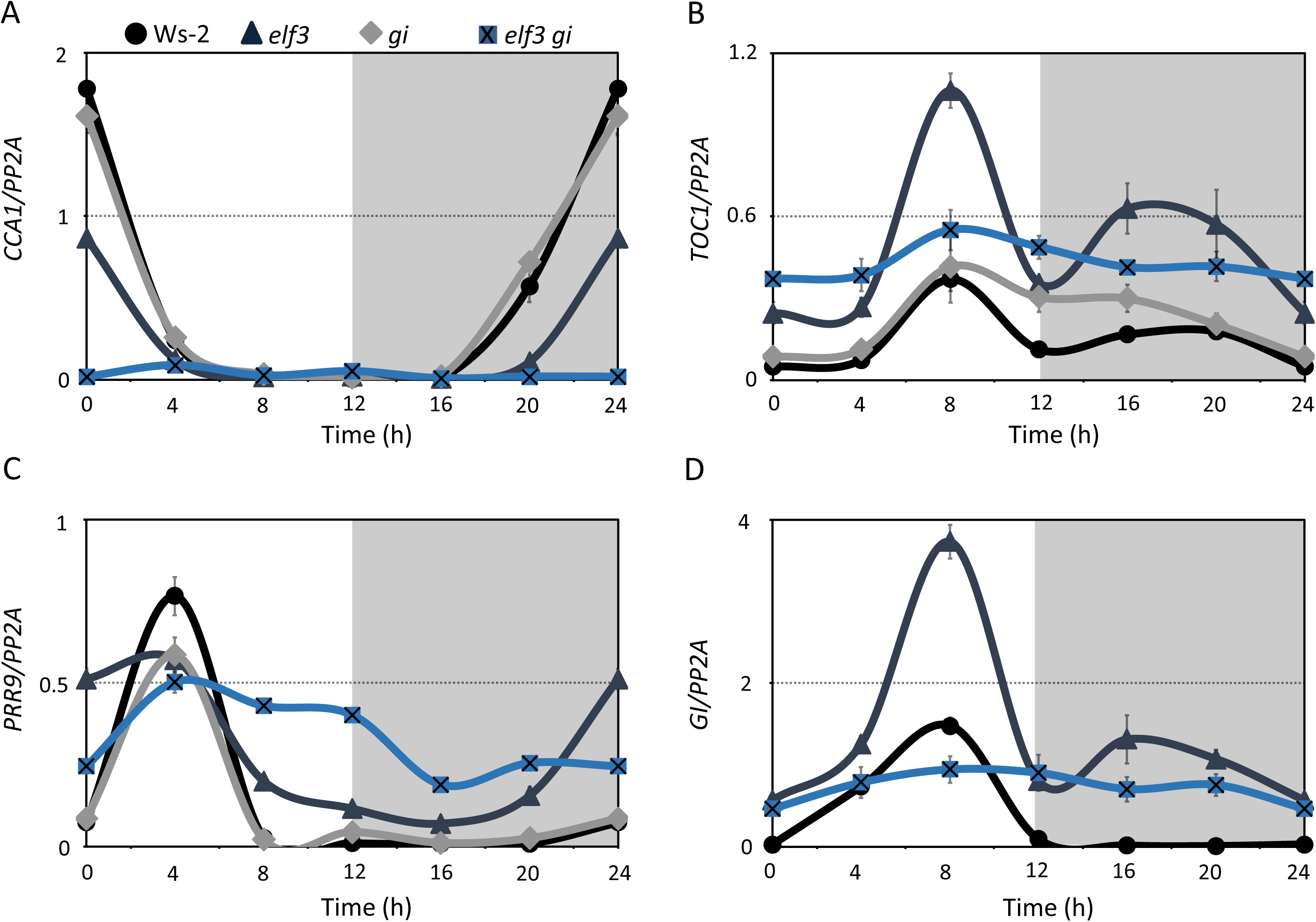
*ELF3* and *GI* are required for clock entrainment. Related to Figure 3. **(A-D)** Transcript accumulation of different circadian clock genes *CCA1* **(A)**, *TOC1* **(B),** *PRR9* **(C)**, and *GI* **(D)** in Ws-2, *elf3*, *gi* and *elf3 gi* (*elf3 gi*) under ND (12h light: 12h darkness). Error bars represent the standard deviation of three technical repeats. Expression levels were normalized to *PROTEIN 19 PHOSPHATASE 2a subunit A3* (*PP2A*). Experiment was repeated with similar results. Non-shaded areas in the graph represent time in LL, shaded areas represent time in DD.

**Figure S4.**
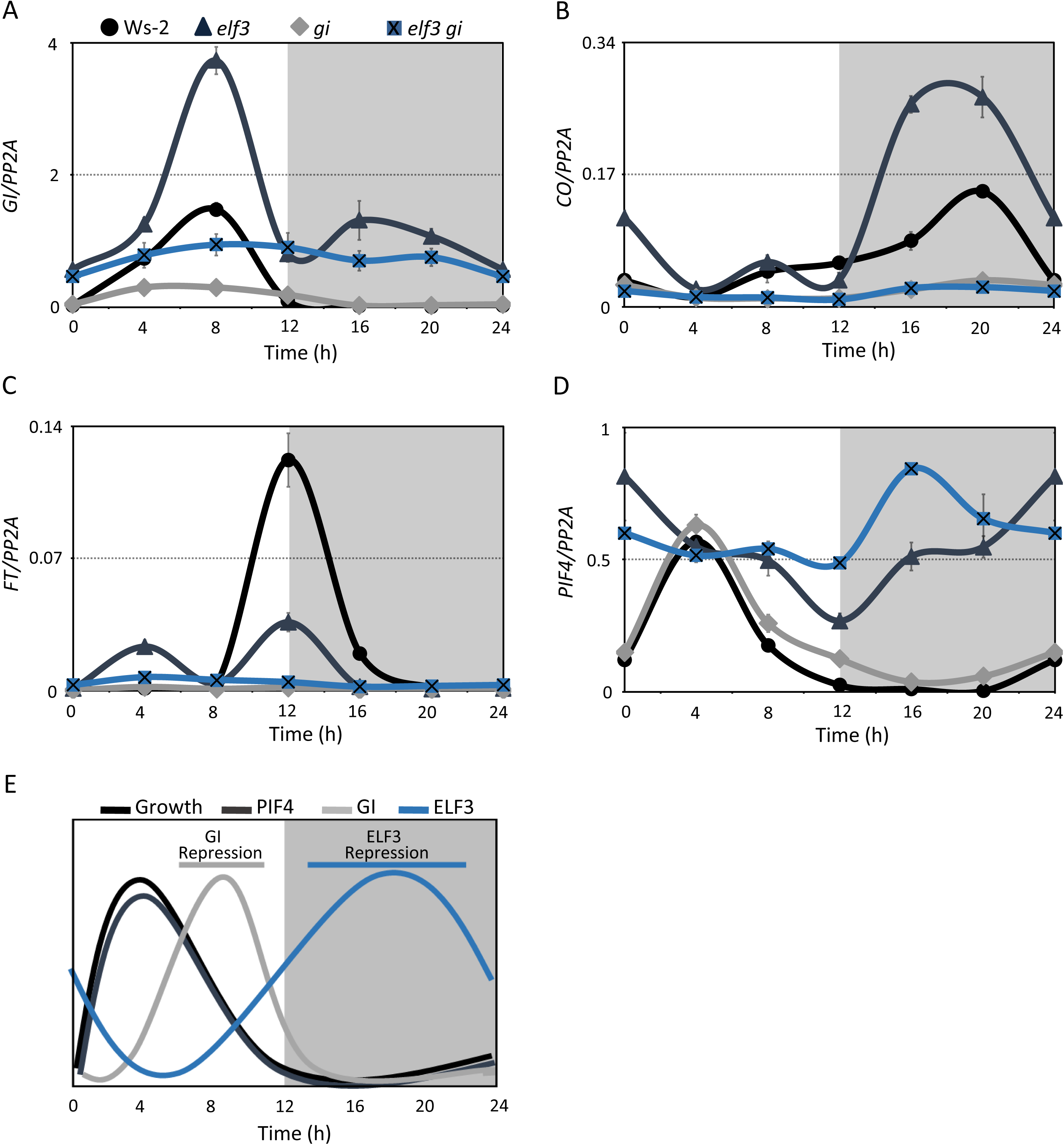
Endogenous and environmental signals synchronization require functional *ELF3* and *GI*. Related to Figure 4. **(A-D)** Transcript accumulation of flowering time genes *GI* **(A)**, *CO* **(B)** and *FT* **(C)** and major growth promoter *PIF4* **(D)** in Ws-2, *elf3*, *gi* and *elf3 gi* (*elf3 gi*) under ND (12h light: 12h darkness). Error bars represent the standard deviation of three technical repeats. Expression levels were normalized to *PROTEIN 19 PHOSPHATASE 2a subunit A3* (*PP2A*). Experiment was repeated with similar results. Non-shaded areas in the graph represent time in LL, shaded areas represent time in DD. **(E)** A model of diurnal control of hypocotyl growth by complementary action of ELF3 and GI. ELF3 and GI repress growth during the night and late-day, respectively by repressing the expression of *PIF4*.

**Figure S5.**
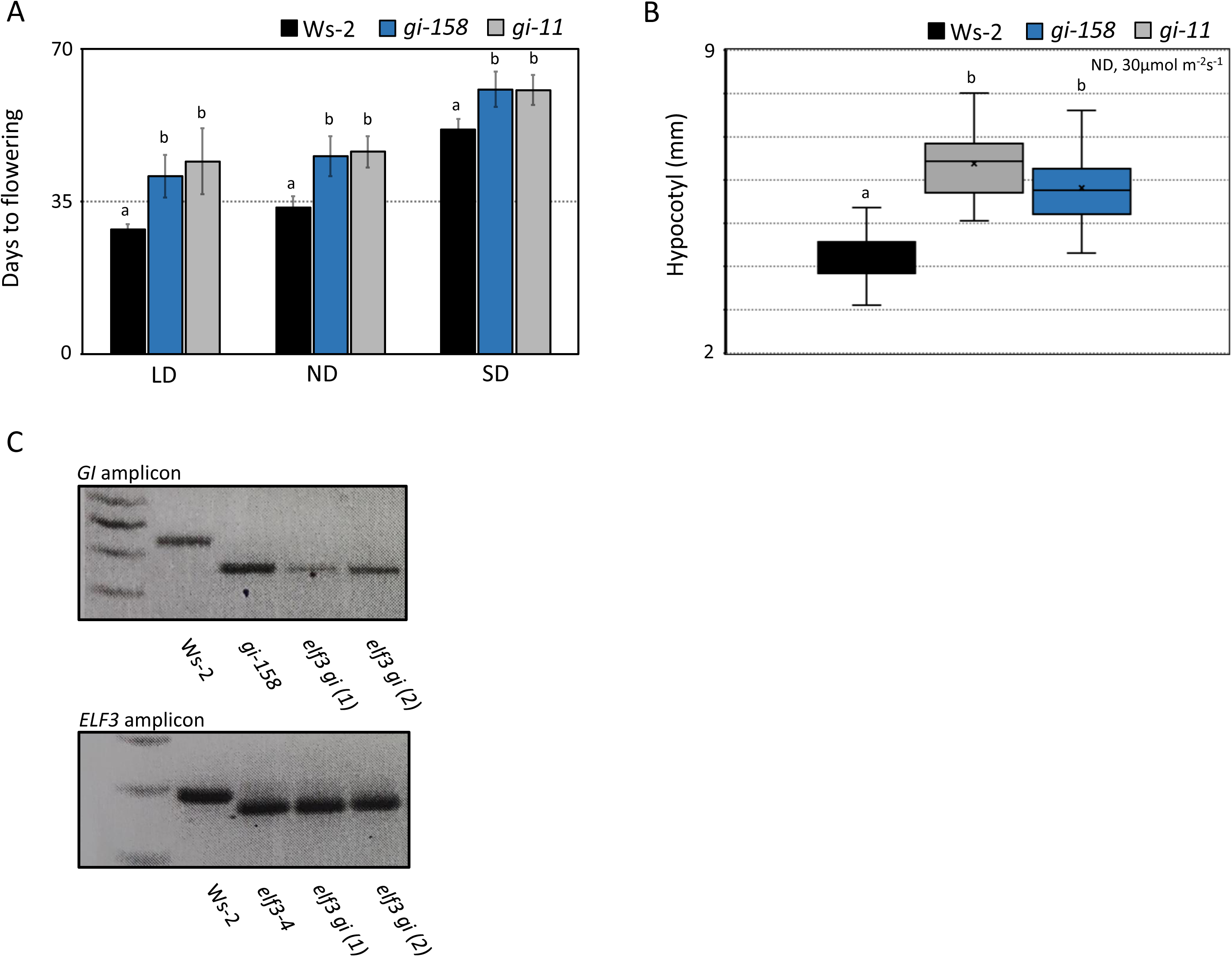
*gi-158* is a null mutant. **(A)** Flowering time of Ws-2, *gi-158* and *gi-11* under LD (16h light: 8 h darkness), 1212 (12h light: 12h darkness), and SD (8h light: 16h darkness). Flowering time was counted as number of days to 1 cm bolt. Error bars represent standard deviation; n≥24. **(B)** Hypocotyl length of Ws-2, *gi-158* and *gi-11* under ND. Significance as described in Figure 1 within a specific photoperiod. **(C)** confirmation of the *elf3 gi* mutant by genotyping. Two independent double mutant lines were obtained after crossing *elf3 gi (1) and elf 3 gi (2)*. After genotypic and phenotypic confirmation, only one line *elf3 gi (1)* was used for further experiments.

**Table S1:** Statistical analysis of the data shown in Figure 1D and S1B-C (ANOVA with post-hoc Tukey HSD Test).

| Light Period | Ws-2 | *elf3* | *gi* | *elf3 gi* |
| --- | --- | --- | --- | --- |
| 00 vs 02 | ** p<0.01 | ** p<0.01 | ** p<0.01 | n.s. |
| 00 vs 04 | ** p<0.01 | ** p<0.01 | ** p<0.01 | ** p<0.01 |
| 00 vs 06 | ** p<0.01 | ** p<0.01 | ** p<0.01 | ** p<0.01 |
| 00 vs 08 | ** p<0.01 | ** p<0.01 | ** p<0.01 | ** p<0.01 |
| 00 vs 10 | ** p<0.01 | ** p<0.01 | ** p<0.01 | ** p<0.01 |
| 00 vs 12 | ** p<0.01 | ** p<0.01 | ** p<0.01 | ** p<0.01 |
| 02 vs 04 | ** p<0.01 | ** p<0.01 | ** p<0.01 | ** p<0.01 |
| 02 vs 06 | ** p<0.01 | ** p<0.01 | ** p<0.01 | ** p<0.01 |
| 02 vs 08 | ** p<0.01 | ** p<0.01 | ** p<0.01 | ** p<0.01 |
| 02 vs 10 | ** p<0.01 | ** p<0.01 | ** p<0.01 | ** p<0.01 |
| 02 vs 12 | ** p<0.01 | ** p<0.01 | ** p<0.01 | ** p<0.01 |
| 04 vs 06 | ** p<0.01 | ** p<0.01 | ** p<0.01 | n.s. |
| 04 vs 08 | ** p<0.01 | ** p<0.01 | ** p<0.01 | ** p<0.01 |
| 04 vs 10 | ** p<0.01 | ** p<0.01 | ** p<0.01 | ** p<0.01 |
| 04 vs 12 | ** p<0.01 | ** p<0.01 | ** p<0.01 | ** p<0.01 |
| 06 vs 08 | ** p<0.01 | * p<0.05 | ** p<0.01 | ** p<0.01 |
| 06 vs 10 | ** p<0.01 | ** p<0.01 | ** p<0.01 | ** p<0.01 |
| 06 vs 12 | ** p<0.01 | ** p<0.01 | ** p<0.01 | ** p<0.01 |
| 08 vs 10 | ** p<0.01 | ** p<0.01 | ** p<0.01 | ** p<0.01 |
| 08 vs 12 | ** p<0.01 | ** p<0.01 | ** p<0.01 | ** p<0.01 |
| 10 vs 12 | n.s. | ** p<0.01 | n.s. | ** p<0.01 |

**Table S2:**
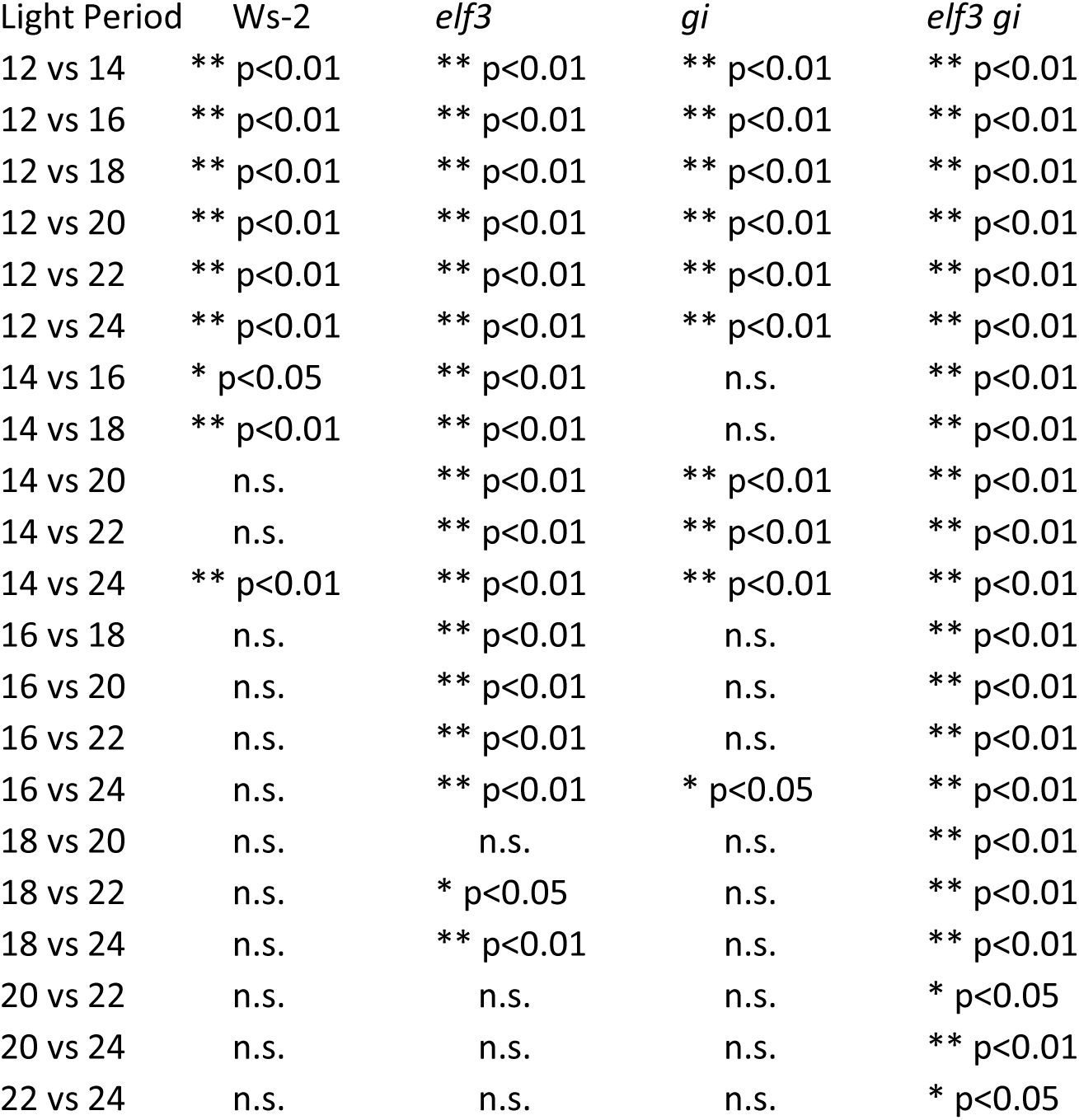
Statistical analysis of the data shown in Figure S1B (ANOVA with post-hoc Tukey HSD Test).

| Light Period | Ws-2 | *elf3* | *gi* | *elf3 gi* |
| --- | --- | --- | --- | --- |
| 12 vs 14 | ** p<0.01 | ** p<0.01 | ** p<0.01 | ** p<0.01 |
| 12 vs 16 | ** p<0.01 | ** p<0.01 | ** p<0.01 | ** p<0.01 |
| 12 vs 18 | ** p<0.01 | ** p<0.01 | ** p<0.01 | ** p<0.01 |
| 12 vs 20 | ** p<0.01 | ** p<0.01 | ** p<0.01 | ** p<0.01 |
| 12 vs 22 | ** p<0.01 | ** p<0.01 | ** p<0.01 | ** p<0.01 |
| 12 vs 24 | ** p<0.01 | ** p<0.01 | ** p<0.01 | ** p<0.01 |
| 14 vs 16 | * p<0.05 | ** p<0.01 | n.s. | ** p<0.01 |
| 14 vs 18 | ** p<0.01 | ** p<0.01 | n.s. | ** p<0.01 |
| 14 vs 20 | n.s. | ** p<0.01 | ** p<0.01 | ** p<0.01 |
| 14 vs 22 | n.s. | ** p<0.01 | ** p<0.01 | ** p<0.01 |
| 14 vs 24 | ** p<0.01 | ** p<0.01 | ** p<0.01 | ** p<0.01 |
| 16 vs 18 | n.s. | ** p<0.01 | n.s. | ** p<0.01 |
| 16 vs 20 | n.s. | ** p<0.01 | n.s. | ** p<0.01 |
| 16 vs 22 | n.s. | ** p<0.01 | n.s. | ** p<0.01 |
| 16 vs 24 | n.s. | ** p<0.01 | * p<0.05 | ** p<0.01 |
| 18 vs 20 | n.s. | n.s. | n.s. | ** p<0.01 |
| 18 vs 22 | n.s. | * p<0.05 | n.s. | ** p<0.01 |
| 18 vs 24 | n.s. | ** p<0.01 | n.s. | ** p<0.01 |
| 20 vs 22 | n.s. | n.s. | n.s. | * p<0.05 |
| 20 vs 24 | n.s. | n.s. | n.s. | ** p<0.01 |
| 22 vs 24 | n.s. | n.s. | n.s. | * p<0.05 |

**Table S3:** Statistical analysis of the data shown in Figure 2E-H (ANOVA with post-hoc Tukey HSD Test).

|  | Time (h) |  |  |  |
| --- | --- | --- | --- | --- |
| LD | 48-56 (1st DD) | 56-72 (1st LL) | 72-80 (2nd DD) | 80-96 (2nd LL) |
| Ws-2 vs *elf3* | ** p<0.01 | n.s. | ** p<0.01 | n.s. |
| Ws-2 vs *gi* | n.s. | * p<0.05 | n.s. | n.s. |
| Ws-2 vs *elf3 gi* | ** p<0.01 | ** p<0.01 | ** p<0.01 | ** p<0.01 |
| *elf3* vs *gi* | ** p<0.01 | n.s. | ** p<0.01 | n.s. |
| *elf3* vs *elf3 gi* | ** p<0.01 | ** p<0.01 | * p<0.05 | ** p<0.01 |
| *gi* vs *elf3 gi* | ** p<0.01 | ** p<0.01 | ** p<0.01 | ** p<0.01 |
| SD | 48-64 (1st DD) | 64-72 (1st LL) | 72-88 (2nd DD) | 88-96 (2nd LL) |
| Ws-2 vs *elf3* | ** p<0.01 | n.s. | ** p<0.01 | n.s. |
| Ws-2 vs *gi* | n.s. | n.s. | n.s. | n.s. |
| Ws-2 vs *elf3 gi* | ** p<0.01 | ** p<0.01 | ** p<0.01 | n.s. |
| *elf3* vs *gi* | ** p<0.01 | * p<0.05 | ** p<0.01 | * p<0.05 |
| *elf3* vs *elf3 gi* | n.s | * p<0.05 | n.s. | n.s. |
| *gi* vs *elf3 gi* | ** p<0.01 | * p<0.05 | ** p<0.01 | * p<0.05 |

